# Assessing the global impact of targeted conservation actions on species abundance

**DOI:** 10.1101/2022.01.14.476374

**Authors:** Sean Jellesmark, Tim M. Blackburn, Shawn Dove, Jonas Geldmann, Piero Visconti, Richard D. Gregory, Louise McRae, Mike Hoffmann

## Abstract

In recent years, vertebrate population abundance has declined at unprecedented rates (WWF 2020). In response, targeted conservation measures – such as breeding programs or species-specific habitat management – have been applied to halt population declines, aid population recovery, and reduce and reverse the loss of biodiversity (Salafsky et al. 2008; Bolam et al. 2020). Until now, assessments of conservation actions have focused on the extent to which they reduce extinction risk, impact populations within protected areas, or increase the global area of land under protection (Hoffmann et al. 2010, 2015; Barnes et al. 2016; Maxwell et al. 2020; Bolam et al. 2020; Grace et al. 2021a). Here, we record and analyze conservation actions for 26,904 vertebrate populations from 4,629 species, to measure the impact of targeted conservation on vertebrate abundance. Using a counterfactual approach to represent population trends in the absence of conservation, we demonstrate that targeted actions have delivered substantial positive effects on the abundance of recipient vertebrate populations worldwide. We show that, in the absence of conservation, a global indicator of vertebrate abundance would have declined even more. Positive population trends were associated with vertebrate populations subject to species or habitat management. We demonstrate that targeted conservation actions can help to reverse global biodiversity loss and show the value of counterfactual analysis for impact evaluation – an important step towards reversing biodiversity declines.

## Introduction

Alterations to global ecosystems have caused widespread declines in biodiversity worldwide (Díaz et al. 2019; WWF 2020), captured by global indicators of the state of biodiversity such as the Red List Index (Butchart et al. 2010), Living Planet Index (LPI) (WWF 2020), and the Biodiversity Intactness Index (Biggs & Scholes 2005). Numerous conservation responses have been implemented to try to halt these declines, from local species-specific efforts, such as ex-situ breeding programs and conservation translocations, to more general large-scale measures, such as the designation of protected areas and international legislation aimed at protecting species (Salafsky et al. 2008; Maxwell et al. 2020; Bolam et al. 2020). Understanding the efficacy of conservation interventions (i.e., to what extent they have contributed to safeguarding biodiversity) is a prerequisite for effective decision-making in conservation and a key priority for researchers, policy and decision makers (Ferraro & Pattanayak 2006; Rose et al. 2018).

Ideally, the impacts of conservation interventions would be assessed using experimental designs that account for potential confounding effects through random allocation of treatment and control groups, such as randomized controlled trials (RCT). If control groups are carefully selected to mimic the treatments in all but the intervention being studied, RCTs offer an experimentally robust approach to estimate the impact of said treatment (Margoluis et al. 2009). However, while experimental designs are possible in certain conservation settings (Wiik et al. 2020), capacity, ethical considerations, and the spatial extent of actions, such as large protected areas, limit the ability to apply experimental evaluations (de Palma et al. 2018; Pynegar et al. 2019). When randomized experiments are not feasible, quasi-experimental designs, based on statistical methods such as matching, can be used instead (Stuart 2010; Joppa & Pfaff 2011; Butsic et al. 2017; Geldmann et al. 2019; Schleicher et al. 2019). For example, annual population counts carried out within and outside protected areas can be matched on observable covariates, using the matched counts to determine how protection relates to population changes (Wauchope et al. 2019a, 2020; Jellesmark et al. 2021).

Alternatively, inferential approaches that use logical arguments (Grace et al. 2021b) or expert knowledge and elicitation to inform counterfactual scenarios can be used to estimate conservation impact (Butchart et al. 2006; Hoffmann et al. 2010; Bolam et al. 2020). For example, Bolam et al (2020) used expert elicitation to estimate the impact of recent conservation actions in averting species extinction and found that, between 1993-2020, conservation may have prevented 21-32 bird and 7-16 mammal extinctions. These studies have advanced the field of counterfactual impact evaluation in conservation science but, until now, we have lacked assessments describing the global impact of conservation actions and responses on species abundance across taxonomic classes.

In this study, we assessed the global impact of species-targeted conservation actions on trends in abundance using population data from the Living Planet Database (LPD)(LPD 2020). Our study has four aims, namely to:

1. Describe targeted conservation actions for species populations in the LPD
2. Assess the impact of conservation actions through a counterfactual approach comparing how population indices differ between conservation targeted and non-targeted populations
3. Measure the impact of conservation on a global population index given different counterfactual scenarios for conservation targeted populations, and
4. Test if specific conservation actions or responses are associated with species’ population trends.

To achieve this, we (1) categorized conservation actions for each managed population in the LPD, (2) created four scenarios to compare trends from conservation targeted populations against matched counterfactuals, (3) created composite population indices in the absence of conservation impact for a subset of populations, and (4) estimated the impact of seven different types of conservation actions on species trends. This allowed us to present a global overview of targeted conservation actions, assess the impact of these actions by approximating how targeted populations were likely to have developed in the absence of conservation, measure the impact of these conservation actions on a global population index, and provide estimates of how each of these conservation actions affects population trends.

## Methods

### The Living Planet Database

The LPD is one of the largest global databases for population time series (WWF 2020). Since 1998, the LPD has provided the vertebrate population abundance data used to estimate the Living Planet Index (Collen et al. 2009; McRae et al. 2017; McRae et al. 2020), one of the key global indicators for biodiversity, and a measure adopted and used by the Convention of Biological Diversity to track progress towards halting the global decline of biodiversity (Butchart et al. 2010; Tittensor et al. 2014; WWF 2020). Today, the LPD is managed by the Zoological Society of London in a collaborative partnership with the World Wildlife Fund, and is continually populated with primary data on vertebrate population abundance, that underpins research in global biodiversity change and is used for indicators to inform both policy makers and the public. The database currently contains information on more than 27,000 populations from almost 5,000 species. These populations are distributed across 11 taxonomic classes (Actinopteri, Coelacanthi, Dipneusti, Elasmobranchii, Holocephali, Myxini, Petromyzonti, Amphibia, Aves, Mammalia, Reptilia). The majority of populations belong to Aves (birds, npops (number of populations) = 10,143), Actinopteri (ray-finned fish, npops = 9,571) and Mammalia (mammals, npops = 5,117) predominantly from North America (npops = 9,692), Europe (npops = 4,997), Latin America and the Caribbean (npops = 4,166) and Asia (npops = 3,835).

Population time-series data are added to the LPD from published or unpublished data if certain data standards are met. First, the data must be for a single species monitored at a defined location over time. Additionally, the species must be a vertebrate (mammal, bird, fish, reptile or amphibian). Several types of abundance data are accepted (Table 1). For example, full population counts are acceptable units of abundance whereas survival rates are not. A minimum of two years of abundance data is required. If a data source contains multiple annual measures, these are converted into a single annual figure using a mean, the peak count, or selecting the most consistently monitored season or month. Besides population data, the source must contain information on the geographic location of the population and the monitoring method. A variety of data sources are accepted given that these are referenced and traceable. This includes peer-reviewed articles from scientific journals, books, reports, online databases and grey literature.

**Table 1.**
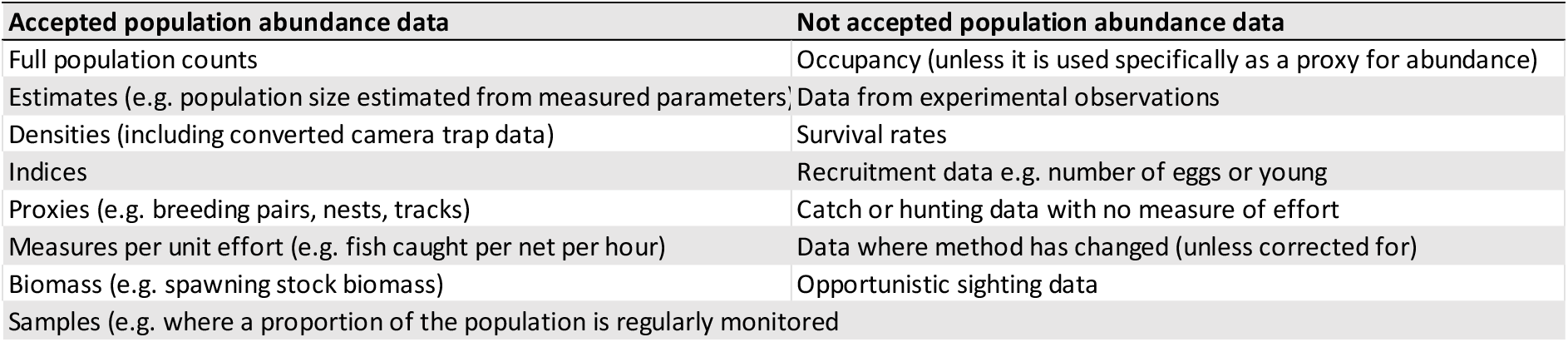
Types of population abundance data that meets the data standards for tracking trends in the abundance of species populations (Accepted) and that does not meet the data standards (Not accepted)

### Database structure

In the LPD, each population is stored with a unique ID and contains annual population data alongside additional information that covers eight broad categories relating to the species or the population. The first category is ‘Base information’, which contains information about the source, the year that the source was published or accessed, the reason for data collection, if the data overlaps with other populations, and the reference for the data. The second category is ‘Taxonomy’. Here, taxonomic information is stored such as the common and Latin species name, Class, Order, Family and Genus. The taxonomic authorities used are:

- *Mammals* – Wilson, D. E. and Reeder, D. M. (2005) Mammal species of the world: a taxonomic and geographic reference (Third Edition). Johns Hopkins University Press, 2,142 pp.
- *Birds* – BirdLife International or IUCN Red List. This is largely consistent with the standard taxonomy for birds (Sibley, C.G. and Monroe, B.L. (1993) Distribution and taxonomy of the birds of the world. Yale University Press: New Haven, USA).
- *Amphibians* – Frost, D. R. (2005) Amphibian Species of the World: an Online Reference (Version 3.0). American Museum of Natural History: New York, USA.
- *Fishes* – Catalog of Fishes researcharchive.calacademy.org/research/ichthyology/catalog/fishcatmain.asp
- *Reptiles* – The Reptiles Database www.reptile-database.org

The third category, ‘Geography’, contains a brief description of a population’s location, latitude, longitude, country and political region. The fourth, ‘Ecology’ category, contains information on the realm and biome in which a population occurs, the habitat type, whether the species is resident in the location, native or alien, and if it is invasive then what impact it has. The fifth category covers population data. Here, population units are recorded along with the sampling method, if the population data have been transformed, the proportion that the population represents of the global population, the annual population value, and if the population increased then additional information about the reason for population increase (reasons can be: Introduction, Recolonisation, Recruitment, Removal of threat, Rural to urban migration, Reintroduction, Range shift, Legal protection, Management, Other, Unknown).

The sixth category contains ‘Protected Area’ information that describes if the population is inside of a protected area, and the type of protected area if so. The seventh category covers ‘Management’ aspects, which indicates if the population is managed, the type of management (see Table 1 for management details), utilized status, CITES and CMS listing. The last category contains ‘Threat’ information, which describes if the population is threatened, the types of threats (threats can be: Habitat Loss, Habitat degradation/Change, Invasive species/genes, Climate change, Pollution, Disease, Exploitation) and whether or not the population is exploited (exploitation can be: Caught and used, Pet trade, Sport hunting, Persecuted as pest, Indirect killing). To ensure consistency, trained personnel record the ‘Management’, ‘Threat’ and ‘Reasons for increase’ information from the original sources using a set of guidelines. This aims to reduce potential bias, but there is likely still to be some individual interpretation of the information in the data source.

### Data selection, management data and coding

We extracted data for every population in the LPD (LPD 2020) – 26,904 populations representing 4,629 species from 11 taxonomic classes. For each of these populations we used the additional data stored in the LPD indicating whether a population was managed, utilised, located inside a protected area, or likely benefitting from conservation action (Table 1). Conservation actions were categorized by extracting all populations in the LPD with management recorded. We first excluded populations where management was unknown (coded as: unmanaged (npops = 9,296); managed (npops = 5,362); or unknown (npops = 12,246). The managed and unmanaged sample contained populations from 182 countries, with 136 countries represented in the managed sample and 169 in the unmanaged sample (Fig. 1). For the populations with management records, we determined if the recorded management qualified as conservation by extracting the management comments from each of the 5,362 managed populations (Table 1). Using these comments on management interventions from the LPD, along with original sources for the population data and additional information provided online (e.g. webpage for specific species recovery projects), we removed populations where the management did not qualify as conservation or research. For the remaining populations we documented conservation and research actions according to the relevant IUCN-CMP conservation actions and research actions classification schemes (Salafsky et al. 2008). These schemes record actions in a hierarchical structure: For example, in the conservation scheme, actions are first divided into seven primary categories (Land & water protection, Land & water management, Species management, Education & awareness, Law & policy, Livelihood Economic & other incentives, External capacity building), and then further divided into detailed sub-categories, such as invasive/problematic species control or species re-introduction (see the IUCN-CMP conservation actions and research actions classification scheme for detailed categories and Supplementary Material for further information and discussion of management information). Populations that could not be categorized in terms of conservation or research actions were removed from the final analysis (npops = 223). Conservation actions were assessed for a total of 14,329 populations in the LPD (53.3% of all), of which 5,243 populations from 1,207 species were recorded as potentially targeted by conservation, and 9,086 populations from 3,106 species were recorded as without targeted conservation. Of these populations, conservation actions were categorized for 4,347 populations, with 41 populations being solely targeted by research actions.

**Figure 1.**
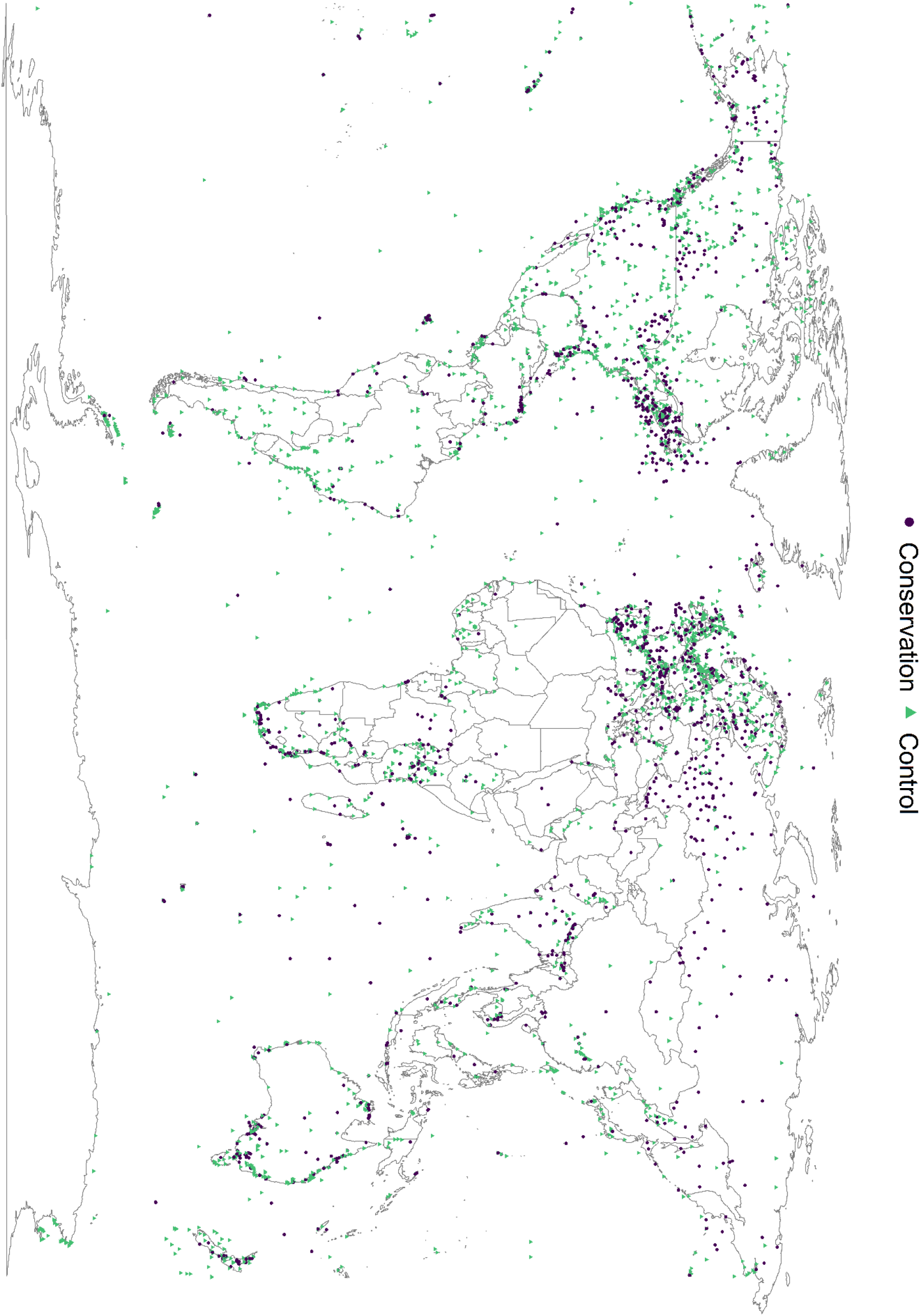
Locations of managed (n = 5,243) and unmanaged populations (n = 9,086).

### Matching populations and calculating trends

To measure the impact of species-targeted conservation actions (hereafter, conservation), we used statistical matching to select the populations used to calculate trends for the populations targeted for conservation and the counterfactual comparison groups, using the MatchIt package (Ho et al. 2021). Matching addresses potential biases between treatment and control populations that could influence the estimates of conservation impacts. This is critical in stochastic environments where the outcome of interest is affected by multiple drivers of change – as is the case for measurements of abundance across ecosystems – and counterfactuals can take many forms (Bull et al. 2020). To characterize this uncertainty, we created four counterfactual scenarios to represent different ways conservation targeted populations could have developed in the absence of conservation.

Scenario one assumed that, in the absence of conservation, populations would have developed similarly to other non-targeted populations from the same genus and political region. Thus, this scenario uses exact matching on genus and region to compare treated populations to non-treated populations from that genus and found in the same region. Exact matching describes similarity between populations using a distance D given a vector X of covariates, where D = 0 for individual i and j if Xi = Xj, and D = ∞ if Xj ≠ Xi. Therefore, each targeted population was matched to all possible control populations with the exact same covariates (Stuart 2010). This creates subcategories based on unique combinations of the selected covariates, assigning populations with similar covariates to the same subcategories. Populations within subcategories that lack either treatment or control are removed. Exact matching allows a single population to have multiple matches, which we use to represent multiple realizations of how a targeted population could have developed.

Scenario two assumed that, in the absence of conservation, a conservation targeted population would show similar trends in abundance to any other population from the same species and country. Relative to the first scenario, exact matching on country and species reduces the sample size but imposes stricter conditions in terms of similarity between the treatment and control populations.

Scenario three assumed that, given no conservation, a targeted population would have developed similarly to a non-targeted population from the same taxonomic class and political region. In addition, we matched each population on the populations’ year of first record and time series length, so that each was matched to a single non-targeted population from the same class and region, and with similar time-series characteristics. We did this using a combination of exact and propensity score matching. Propensity score matching uses logistic regression to predict a probability of receiving treatment, which in this case is targeted conservation, given a set of observed predictors (Williamson & Forbes 2014). Taxonomic class and global region were included as exact matches, whereas the first year of observation and time series length were included as continuous covariates using one-to-one covariate matching without replacement based on the propensity score. This ensured that the targeted sample and the counterfactual contained the same number of populations, each population compared to its closest match given the observed covariates. Including the year of the time-series was to address any overarching changes within the regions that might mean that comparing populations from different periods of time would be problematic.

In the fourth scenario, we made no assumptions about how the targeted populations would have developed without conservation. We thus compared the full sample of conservation targeted populations for the conservation group with the full sample of populations without conservation as the counterfactual group.

For each of the four scenarios, we created multi-species indices of relative abundance using the rlpi package (Freeman et al. 2020). Here, annual population growth rates are modelled using the chain method for populations with fewer than six data points, and a Generalized Additive Model (GAM) for populations with six or more observations (Collen et al. 2009; McRae et al. 2017). For species with multiple populations, the estimated annual trends were averaged into a single annual trend following 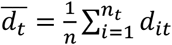 where *n_t_* is the number of populations and *d_it_* is the annual population change rate in a given year. The rate of change is given by 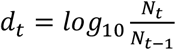 where *N* is the population estimate and *t* is the year. Annual log growth rates were capped between −1 and 1. Indices were created based on a geometric mean approach using the log-transformed growth rates where the index year *I*_0_ = 1 and the following indices are calculated as 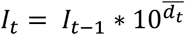 (McRae et al. 2017). The 95% confidence intervals were generated with 10,000 bootstrap replicates across species-level annual growth rates (Collen et al. 2009).

Our approach is similar to that used to create the LPI, except that no taxonomic or other weighting was applied, so that each species was weighted equally. The LPI aims to characterize global trends in vertebrate populations in a balanced fashion, and therefore applies weighting to account for taxonomic and geographical inequalities in the sampled data (McRae et al. 2017). However, we set out to estimate the impact of conservation actions on target populations using a matching approach which similarly corrects for bias. Furthermore, we created the four counterfactuals from matched subsamples of the LPD. The weighted approach is not suitable because these samples are much smaller than the actual LPD, and not randomly selected. Therefore, we did not apply the LPI weighting, as this could potentially exacerbate the effect of any selection biases in ways that would be difficult to interpret.

Clusters of populations with extreme abundance changes and time series length have been shown to influence population trend indicators (Wauchope et al. 2019b; Leung et al. 2020). We therefore tested the sensitivity of our indices by recreating them without the top and bottom 1 % quantiles of species with increasing and decreasing populations, and by restricting populations to those with time series spanning a minimum of 5 and 10 years.

#### Calculating the impact of conservation on a global index of species abundance

To assess the wider global impact of conservation on populations, we used a quantitative method that built on Hoffmann et al (2010). First, we calculated a global vertebrate population index using all trend data in the LPD. Then, to evaluate how conservation actions have affected global species abundance, we calculated the index under alternative assumptions into three counterfactual indices. The first was a simple population index excluding all conservation targeted populations. This index was calculated by removing the populations with records of targeted conservation but otherwise using all the available LPD population data.

We calculated a second population index where the impact of targeted conservation actions was excluded by assuming stable trends for otherwise increasing populations with records of conservation actions. For this index, we first identified those populations exhibiting an observed increase over the time series, and then selected all populations categorized as ‘conservation targeted’ and for which information about the reason for population increase was recorded. From this sample, we selected increasing populations with a plausible link between the observed change and a conservation intervention (see Supplementary Material for the full method of validating this link). The reasons for population increase that we selected are underlined in Table 2. For this subsample, we replaced the observed abundance estimates with a constant, thus assuming that trends for these populations would have at least remained stable without conservation. The index was then calculated from all of the available LPD population data, but with constant annual abundance estimates for the selected subset of populations.

**Table 2.** Table of the information extracted from the LPD on the variables used for categorizing conservation actions and analysis. The reasons for population increases that have been used to calculate the global impact of conservation on the unweighted LPI are underlined.

| Variable | Description |
| --- | --- |
| **Species** | Taxonomic information according to the latest authority for that species |
| **Genus** | Taxonomic information according to the latest authority for that genus |
| **Class** | Taxonomic information according to the latest authority for that class. Coding: 'Actinopteri', 'Coelacanthi', 'Dipneusti', 'Elasmobranchii', 'Holocephali', 'Myxini', 'Petromyzonti', 'Amphibia', 'Aves', 'Mammalia', 'Reptilia'. |
| **Country** | The country (or countries) that the population occurs in from the list. Marine data are allocated a country if it is within its EEZ, or as International Waters. Multiple countries are selected in order of proportion of the population it represents starting with the greatest. |
| **Region** | The political region a country is assigned to. Coding: 'Africa', 'Antarctic', 'Asia', 'Europe', 'International Waters', 'Latin America and Caribbean', 'North America', 'Oceania' |
| **First year of observation** | The first recorded year with an abundance estimate for a given population |
| **Time series length** | The number of years from first to last population abundance estimate |
| **Managed** | A population that receives targeted management (some of which involves sustainable use). This is usually to promote recovery in a population or can incentivise its use for conservation. It can include measures to stem 'unsustainable' population growth. Coding: 'Yes', 'No', 'Unknown' |
| **Utilised** | A population that is intentionally regularly or systematically utilised, either individuals or eggs. This may be sustainable or unsustainable, and the population does not necessarily have to be threatened by use or overexploited. This refers to consumptive use whereby individuals or parts of individuals are removed from the wild. Coding: 'Yes', 'No', 'Unknown' |
| **Targeted conservation actions** | A population that is intentionally targeted by conservation. Coding of conservation actions follows categories in Salafsky et al 2008 |
| **Reason for population increase** | Indicates the reasons given by the data source for any increase in the population. Coding: '<u>Introduction</u>', '<u>Recolonisation</u>', '<u>Recruitment</u>', '<u>Removal of threat</u>', 'Rural to urban migration', '<u>Reintroduction</u>', '<u>Range shift</u>', '<u>Legal protection</u>', '<u>Management</u>', 'Other', 'Unknown'. |
| **Protected area status** | Indicates if the population is inside a protected area. Coding: 'Both', 'No', 'No (area surrounding PA)', 'No (Large survey area)', 'Unknown', 'Yes'. |

Finally, we calculated a population index where the impacts of targeted and collateral conservation were excluded (Hoffmann et al. 2015). This population index, excluding the impact of both targeted and collateral conservation, was calculated similar to the second index but in addition excluding the impact of collateral conservation. By collateral, we mean that a population could have benefitted from conservation without being specifically targeted, which we defined as having increasing population trends within a protected area, while not being specifically chosen for any of the targeted actions. This index was therefore calculated using the full LPD data but assuming stable trends for the same populations as in the second index and additionally populations without targeted conservation, but which were inside a protected area and had a reason for population increase recorded. Population indices were calculated using the rlpi package (Freeman et al. 2020) without applying taxonomic or geographical weighting. To visualize the impact of conservation, we plotted the difference between the reference population index, calculated using all of the population trend data in the LPD, and the three potential scenarios that represent the reference index without the likely impact of conservation.

#### Mixed model

We compared the effects of the seven primary targeted conservation actions on abundance trends (the log of the summed rate of population change) using a mixed model framework (McRae et al. 2020). The rlpi package generates a matrix of annual rates of population change for each population. We summed these rates into a logged value of total change for each population. To test the effects of conservation actions in general, and of the different types of interventions, two separate models were specified.

In the first, conservation actions were aggregated into a single fixed effect binary variable (1 = targeted or 0 = not targeted by conservation) to estimate the overall effect of actions regardless of action type.

In the second, we specified a binary variable for each of the seven main conservation actions. This allowed us to estimate the effect of conservation actions (model one) and then disentangle the individual effects of specific types of actions (model two). We included time series length, taxonomic class and the utilization status of each population as fixed effects, as these have been shown to affect abundance changes (Wauchope et al. 2019b; McRae et al. 2020). We specified similar random effect structures as in McRae et al (2020), including Family, Genus and population location to account for taxonomic and site specific effects (Model one: sum_lambda ~ 0 + TS_length + Utilised + Conservation + Class + (1|Family/Binomial) + (1|Location); Model two: lambda_sum ~ 0 + Utilised + ts_length + land_water_protection + land_water_management + species_management + education_awareness + law_policy + incentives + external_capacity + research + Class + (1|Family/Binomial) + (1|Location)).

## Results

### Conservation actions in the LPD

Mammals had the highest number of managed populations in the LPD, albeit across a relatively low number of species (nspp (number of species) = 244, npops = 2,200), followed by fish (nspp = 548, npops = 1,994), birds (nspp = 305, npops = 756), reptiles (nspp = 58, npops = 220) and amphibians (nspp = 52, npops = 73). The taxonomic classes included in the fish, mammal and bird groups, maps of the starting year (Fig. S2) and the length (Fig. S3) of the population time series, are all given in the Supplementary Material.

Species management was the most frequent conservation action (n = 2,937), followed by land & water management (n = 1,095), and law & policy actions (n = 467) (Fig. 2; see Fig. S1 for detailed categories). Conservation actions differed between classes, with species management being the most abundant action for mammals and fishes, while land & water management was more frequently applied for birds.

**Figure 2.**
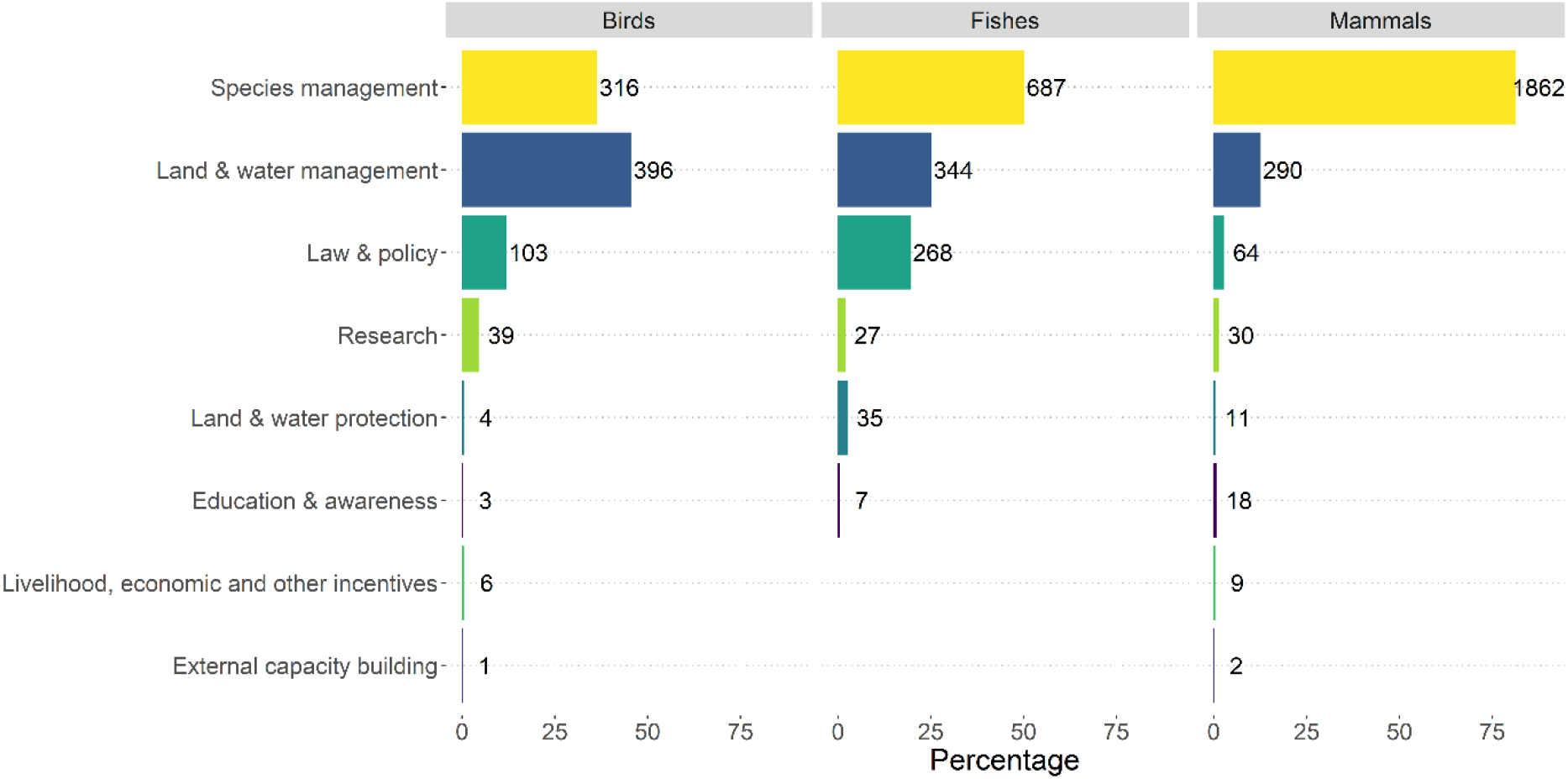
Number of targeted populations and the relative percentage of conservation actions for fish, birds and mammals. For each of the three groups with targeted conservation actions, the x-axis shows the percentage of populations targeted by the seven primary conservation actions and research (Salafsky et al. 2008). The number of targeted populations is shown for each bar.

### Impact of conservation under four different counterfactual scenarios

Trends for populations targeted by conservation actions increased consistently and strongly when compared with counterfactuals (Fig. 3). The largest difference was observed when comparing populations of the same species within the same country (scenario 2). Here, the index for the conservation targeted sample increased from 1 to 3.36 (234% increase), whereas the counterfactual sample increased to just 1.01 (1% increase). The smallest difference between the indices was observed in scenario 3 (matching on taxonomic class, region, time series length and starting year) where the index for the conservation targeted sample increased to 1.6 (60% increase) while the counterfactual decreased to 0.79 (21% decrease). Sensitivity tests showed that the conservation targeted population indices remained higher than the counterfactual in all cases (Fig. S4. See Fig. S5 for the number of species in each class within each scenario).

**Figure 3.**
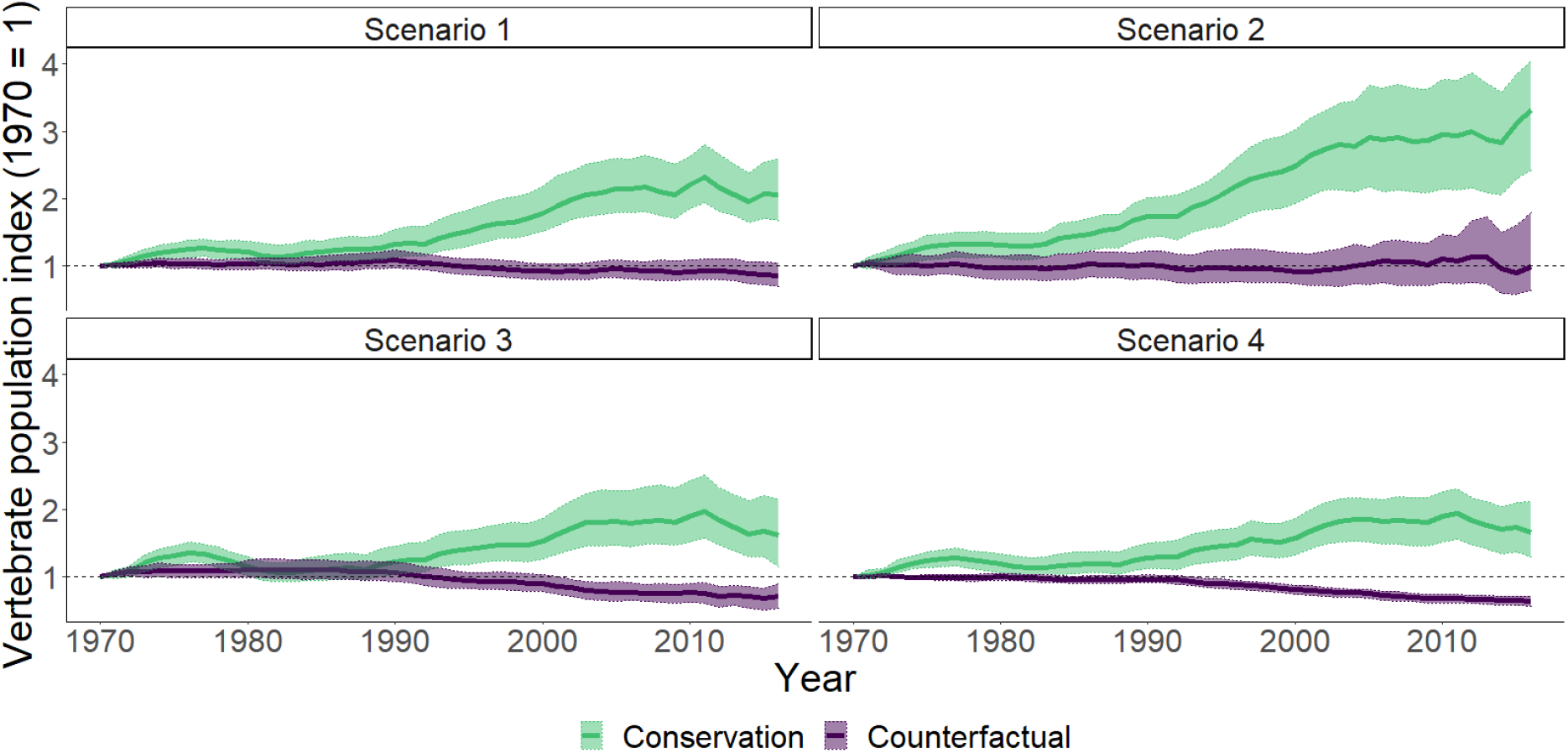
Vertebrate population trends for species subject to conservation actions or responses (in green – upper lines) and not targeted by conservation responses (in purple - lower lines) representing counterfactual species trends. Shaded areas show 95% confidence intervals. Dashed line equals index 1. Scenario 1 – Genus + Region; Conservation + 103%; nspp = 785, npop = 3,377, Counterfactual - 30%; nspp = 1,001, npop = 3,463: Scenario 2 – Species + Country; Conservation + 234%; nspp = 348, npop = 1483,Counterfactual + 1%; nspp = 347, npop = 895: Scenario 3 – Class + Region + Time series length + Start year; Conservation + 60%; nspp = 1,010, npop = 2,929, Counterfactual - 21%; nspp = 1,449, npop = 2,929: Scenario 4 – Full sample; Conservation + 75%; nspp = 1,207, npop = 5,243, Counterfactual - 35%; nspp = 3,099, npop = 9,071

### Impact of conservation on global species abundance

Our global vertebrate population index decreased by 24% between 1970-2016 but would likely have declined by 31% (7% points more) if conservation targeted populations had remained stable, or by 32% (8% points more) if both conservation targeted populations and populations affected by collateral conservation remained stable (Fig 4, Fig S7).

**Figure 4.**
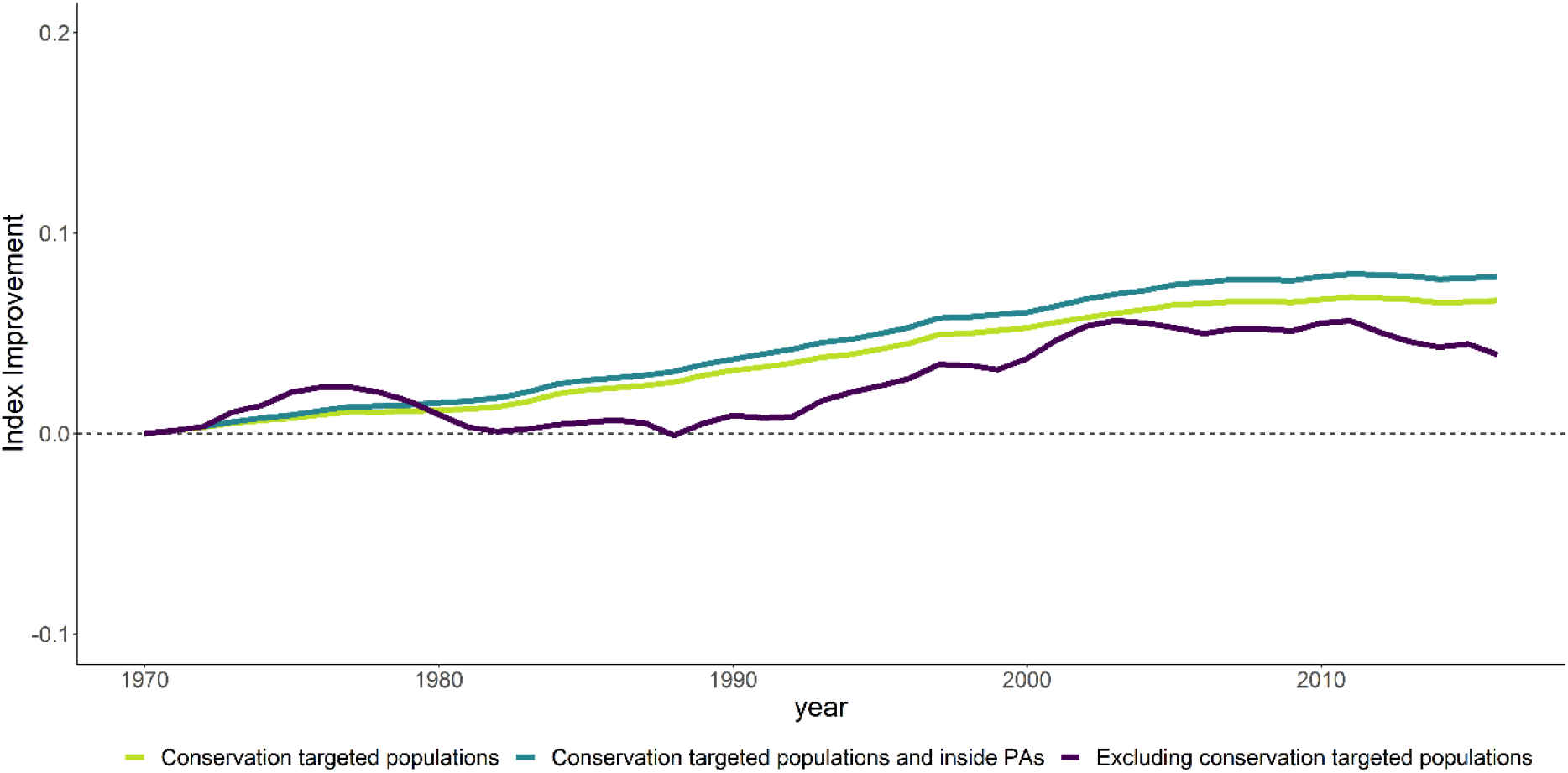
Improvements in invertebrate population trends when assuming stable trends for populations with increasing population trends attributable to conservation (light green – middle line), when assuming stable trends for increasing conservation targeted populations and populations inside PAs attributable to conservation (turquoise – top line) and when excluding populations targeted by conservation actions (purple – bottom line). Improvements are calculated by subtracting each of the three counterfactual population trends from the global reference trend. The dashed line represents no difference between the reference trend and any of the three alternative trends. See Fig. S7 for the original trends.

### Drivers of population trends estimated from mixed models

Conservation actions had a positive effect on targeted populations (Estimate = 0.12, Std Error = 0.02, t value = 5.8). Land & water management, species management and land & water protection actions for species were highly associated with population increases, suggesting a particularly strong effect of actions within these three conservation categories (Fig. 5). Research actions were negatively associated with population trends. Longer population time series were more likely to have increased, while utilized populations did not display a clear trend.

**Figure 5.**
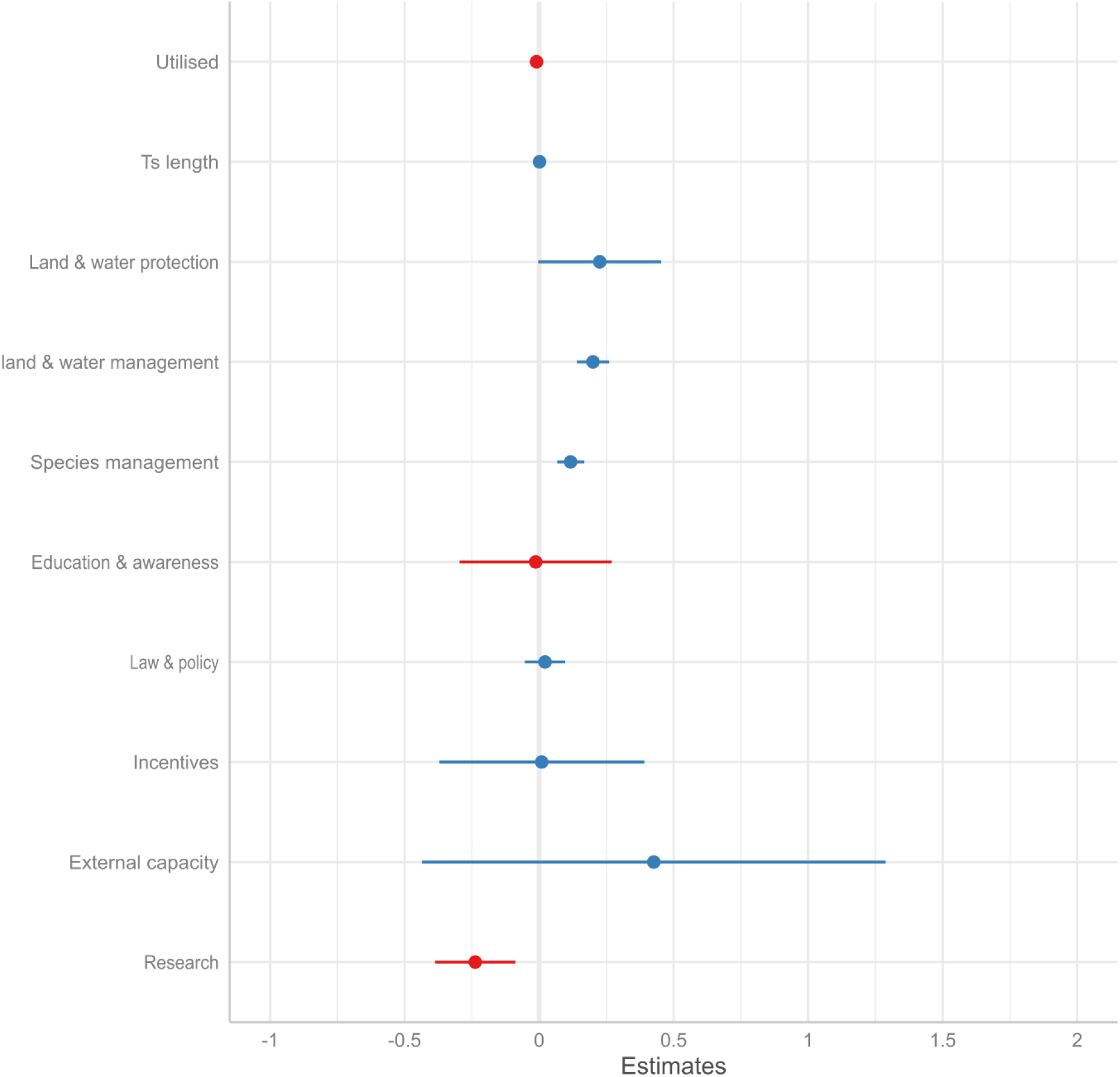
Parameter estimates (estimated total change on the log scale) for the seven primary types of conservation actions, research, utilization status and time series length. See Table S1 for parameter estimates, standard errors and t values.

## Discussion

Our analyses, using one of the largest global datasets of population time-series, show that conservation actions have had a positive influence on global vertebrate populations. Our results were consistent and robust, with a positive impact of conservation detected for all scenarios of counterfactual population developments tested, substantiated by a marked difference between global indices including and excluding the impact of conservation. Furthermore, we saw an effect of conservation not only when comparing the relative difference between treatments and counterfactuals: increasing population trends in absolute terms were also more likely for populations targeted by conservation actions, especially land & water management and species management. Our findings therefore suggest that conservation has delivered substantial benefits to targeted populations.

Our analysis demonstrates the importance of conservation actions that are less frequently evaluated, and thus expands on previous large-scale evaluation efforts within conservation science. Previous efforts have, to a large degree, focused on protected areas (Geldmann et al. 2013a, 2013b, 2019; Butchart et al. 2015; Barnes et al. 2016; Wauchope et al. 2019a; Cazalis et al. 2020, 2021; Maxwell et al. 2020; Terraube et al. 2020). However, protected areas are under a wide variety of management regimes, with large differences in management effectiveness (Geldmann et al. 2015). This means that the percentage of area protected by itself is inadequate to measure conservation effectiveness (Visconti et al. 2019; Rodrigues & Cazalis 2020), and potentially conceals the effectiveness of certain conservation interventions. By identifying the effects of specific conservation actions targeting populations of vertebrate species, we demonstrate the positive impact of conservation efforts on vertebrate populations globally.

Conservation actions without any immediate effect on population trends, such as education & awareness (Fig. 5) can still provide important conservation benefits. We assessed the impact of conservation on population abundance, but there are many alternative outcome metrics which could have illustrated the effectiveness of these actions. For example, we found no effect of awareness & education as this category of conservation might not cause population increases directly. Instead, awareness & education can work indirectly by providing funding for conservation organizations and by giving mandate and support for legal protection to governments.

Accurate assessments of conservation impact depend on accessible and representative data across different aspects of biodiversity. Likewise, appropriately specified counterfactuals require information about the covariates that affect the sample. Currently, global biodiversity data are taxonomically and geographically biased (McRae et al. 2017; Troudet et al. 2017; Hughes et al. 2021). Additionally, fine-scale data and contextual information for conservation targeted populations, such as the type or duration of conservation management, are extremely limited. For example, we could not distinguish between population trends pre and post intervention, as such data do not readily exist on a global level. Advances in remote sensing are already reducing this knowledge gap somewhat, with data derived from remote sensing images widely used to inform where to target conservation actions and evaluate the impact of environmental policy (Chen et al. 2019; Runting et al. 2020). However, many measures cannot be proxied by land cover (for example reintroduction programmes) and must therefore be complemented by in-situ monitoring. Here, we show the relevance of retaining and standardizing such local information, but also that widespread systematic monitoring is required to improve evaluation efforts, especially outcome-based assessments, and to determine the progress towards future biodiversity targets. For example, future monitoring records could be standardized to capture when monitored populations were targeted by conservation actions, and the associated costs. Additional information, such as the temporal exposure to a conservation intervention, would allow effect estimates to be derived with greater confidence from more reliable study designs (de Palma et al. 2018; Christie et al. 2019; Wauchope et al. 2020).

Our analysis calculating global indices with and without the impact of conservation likely underestimates the impact of conservation. First, we assumed that, in the absence of conservation, conservation targeted populations that increased would have remained stable. This is in stark contrast to the general pattern of global declines (WWF 2020). Second, because of data limitations, only a small subset of the total LPD has a reason for increase recorded. Furthermore, effective conservation is not conditional on population increase. Instead, for conservation to be effective requires only that the outcome of interest is improved by conservation, relative to a scenario without conservation.

While populations in the LPD only represent a fraction of global biodiversity, our results offer a glimmer of hope and underpin the importance of conservation efforts in halting the global loss of biodiversity. As the global parties to the Convention on Biological Diversity are preparing the Post-2020 Global Biodiversity Framework, it is relevant to reflect on conservation progress made over the last decade, as well as how to measure progress towards achieving these targets. One of the most prominent elements of the CBDs Strategic Plan for biodiversity 2011-2020 was focused on the establishment of effective and representative networks of protected areas (Sustainable Development Goals – Aichi Target 11), and post-2020 targets set out more ambitious targets in this regard, currently suggesting. that 30% of global terrestrial area should be placed under formal protection (First draft of the post-2020 global biodiversity framework). It is important to recognize the progress made towards increasing the global coverage of protected area, with marked increases observed both on land, in freshwater environments and in the marine realm (Maxwell et al. 2020; UNEP-WCMC & IUCN 2021). However, vertebrate populations continue to decline (WWF 2020). Similarly, without increased conservation efforts, global biodiversity projections predict continuous declines in the future (Leclère et al. 2020), highlighting the need for effective conservation actions and outcome-based targets (Butchart et al. 2015; Visconti et al. 2019; Pressey et al. 2021). We show that such targeted conservation interventions can be highly effective.

## References

Barnes MD et al. 2016. Wildlife population trends in protected areas predicted by national socio-economic metrics and body size. Nature Communications 7:1–9. Nature Publishing Group.

Biggs R, Scholes RJ. 2005. A biodiversity intactness index. Nature 434:45–49. Available from http://go.galegroup.com/ps/i.do?id=GALE%7CA185471773&v=2.1&u=ntu&it=r&p=AONE&sw=w.

Bolam FC et al. 2020. How many bird and mammal extinctions has recent conservation action prevented? Conservation Letters:e12762. Available from http://doi.wiley.com/10.1111/conl.12762.

Bull JW, Strange N, Smith RJ, Gordon A. 2020. Reconciling multiple counterfactuals when evaluating biodiversity conservation impact in social-ecological systems. Conservation Biology:cobi.13570. Available from https://onlinelibrary.wiley.com/doi/abs/10.1111/cobi.13570.

Butchart SHM et al. 2010. Global biodiversity: Indicators of recent declines. Science 328:1164–1168.

Butchart SHM et al. 2015. Shortfalls and Solutions for Meeting National and Global Conservation Area Targets. Conservation Letters 8:329–337.

Butchart SHM, Stattersfield AJ, Collar NJ. 2006. How many bird extinctions have we prevented? ORYX DOI: 10.1017/S0030605306000950.

Butsic V, Lewis DJ, Radeloff VC, Baumann M, Kuemmerle T. 2017. Quasi-experimental methods enable stronger inferences from observational data in ecology. Basic and Applied Ecology 19:1–10. Elsevier GmbH. Available from http://dx.doi.org/10.1016/j.baae.2017.01.005.

Cazalis V, Barnes MD, Johnston A, Watson JEM, Şekercioğlu CH, Rodrigues ASL. 2021. Mismatch between bird species sensitivity and the protection of intact habitats across the Americas. Ecology Letters:ele.13859. Available from https://onlinelibrary.wiley.com/doi/10.1111/ele.13859.

Cazalis V, Princé K, Mihoub JB, Kelly J, Butchart SHM, Rodrigues ASL. 2020. Effectiveness of protected areas in conserving tropical forest birds. Nature Communications 11:1–8. Springer US. Available from http://dx.doi.org/10.1038/s41467-020-18230-0.

Chen C et al. 2019. China and India lead in greening of the world through land-use management. Nature Sustainability 2:122–129. Springer US. Available from http://dx.doi.org/10.1038/s41893-019-0220-7.

Christie AP, Amano T, Martin PA, Shackelford GE, Simmons BI, Sutherland WJ. 2019. Simple study designs in ecology produce inaccurate estimates of biodiversity responses. Journal of Applied Ecology:2742–2754.

Collen B, Loh J, Whitmee S, McRae L, Amin R, Baillie JEM. 2009. Monitoring change in vertebrate abundance: the living planet index. Conservation biology : the journal of the Society for Conservation Biology 23:317–27. Available from http://doi.wiley.com/10.1111/j.1523-1739.2008.01117.x.

de Palma A, Sanchez-Ortiz K, Martin PA, Chadwick A, Gilbert G, Bates AE, Börger L, Contu S, Hill SLL, Purvis A. 2018. Challenges With Inferring How Land-Use Affects Terrestrial Biodiversity: Study Design, Time, Space and Synthesis. Page Advances in Ecological Research, 1st edition. Elsevier Ltd. Available from http://dx.doi.org/10.1016/bs.aecr.2017.12.004.

Díaz S et al. 2019. Pervasive human-driven decline of life on Earth points to the need for transformative change. Science 366.

Ferraro PJ. 2009. Counterfactual thinking and impact evaluation in environmental policy. New Directions for Evaluation 2009:75–84. Available from https://onlinelibrary.wiley.com/doi/10.1002/ev.297.

Ferraro PJ, Pattanayak SK. 2006. Money for Nothing? A Call for Empirical Evaluation of Biodiversity Conservation Investments. Plos biology 4:482–488.

Freeman R, McRae L, Deinet S, Amin R, Collen B. 2020. rlpi: Tools for calculating indices using the Living Planet Index method. Available from https://github.com/Zoological-Society-of-London/living_planet_index.

Geldmann J et al. 2015. Changes in protected area management effectiveness over time: A global analysis. Biological Conservation 191:692–699. Elsevier B.V. Available from http://dx.doi.org/10.1016/j.biocon.2015.08.029.

Geldmann J, Barnes M, Coad L, Craigie ID, Hockings M, Burgess ND. 2013a. Effectiveness of terrestrial protected areas in reducing habitat loss and population declines. Biological Conservation 161:230–238. Elsevier Ltd. Available from http://dx.doi.org/10.1016/j.biocon.2013.02.018.

Geldmann J, Barnes M, Coad L, Craigie ID, Hockings M, Burgess ND. 2013b. Effectiveness of terrestrial protected areas in reducing habitat loss and population declines. Biological Conservation 161:230–238. Available from https://linkinghub.elsevier.com/retrieve/pii/S0006320713000670.

Geldmann J, Manica A, Burgess ND, Coad L, Balmford A. 2019. A global-level assessment of the effectiveness of protected areas at resisting anthropogenic pressures. Proceedings of the National Academy of Sciences of the United States of America 116:23209–23215. Available from https://doi.org/10.1073/pnas.1908221116.

Grace MK et al. 2021a. Testing a global standard for quantifying species recovery and assessing conservation impact. Conservation Biology:1–17.

Grace MK et al. 2021b. Building robust, practicable counterfactuals and scenarios to evaluate the impact of species conservation interventions using inferential approaches. Biological Conservation 261:109259. Elsevier Ltd. Available from https://doi.org/10.1016/j.biocon.2021.109259.

Ho D, Imai K, King G, Stuart E, Whitworth A, Greifer N. 2021. Package ‘ MatchIt.’ Available from https://github.com/kosukeimai/MatchIt.

Hoffmann M et al. 2010. The impact of conservation on the status of the world’s vertebrates. Science 330:1503–1509.

Hoffmann M, Duckworth JW, Holmes K, Mallon DP, Rodrigues ASL, Stuart SN. 2015. The difference conservation makes to extinction risk of the world’s ungulates. Conservation Biology 29:1303–1313.

Hughes AC, Orr MC, Ma K, Costello MJ, Waller J, Provoost P, Yang Q, Zhu C, Qiao H. 2021. Sampling biases shape our view of the natural world. Ecography 44:1259–1269.

Jellesmark S, Ausden M, Blackburn TM, Gregory RD, Hoffmann M, Massimino D, McRae L, Visconti P. 2021. A counterfactual approach to measure the impact of wet grassland conservation on UK breeding bird populations. Conservation Biology:cobi.13692. Available from https://onlinelibrary.wiley.com/doi/10.1111/cobi.13692.

Joppa LN, Pfaff A. 2011. Global protected area impacts. Proceedings of the Royal Society B: Biological Sciences 278:1633–1638. Available from https://royalsocietypublishing.org/doi/10.1098/rspb.2010.1713.

Leclère D et al. 2020. Bending the curve of terrestrial biodiversity needs an integrated strategy. Nature 2018. Available from https://doi.org/10.1038/s41586-020-2705-y%0Ahttp://www.nature.com/articles/s41586-020-2705-y.

Leung B, Hargreaves AL, Greenberg DA, Mcgill B, Dornelas M, Freeman R. 2020. Clustered versus catastrophic global vertebrate declines. Nature DOI: 10.1038/s41586-020-2920-6. Springer US. Available from http://dx.doi.org/10.1038/s41586-020-2920-6.

LPD. 2020. Living Planet Index database. Available from https://livingplanetindex.org (accessed November 16, 2020).

Margoluis R, Stem C, Salafsky N, Brown M. 2009. Design alternatives for evaluating the impact of conservation projects. New Directions for Evaluation 2009:85–96. Available from http://doi.wiley.com/10.1002/ev.298.

Maxwell SL et al. 2020. Area-based conservation in the twenty-first century. Nature 586:217–227. Springer US. Available from http://dx.doi.org/10.1038/s41586-020-2773-z.

McRae L, Deinet S, Freeman R. 2017. The diversity-weighted living planet index: Controlling for taxonomic bias in a global biodiversity indicator. PLoS ONE 12.

McRae L, Freeman R, Geldmann J, Moss GB, Kjær-hansen L, Burgess D. 2020. A global indicator of utilised wildlife populations : regional trends and the impact of management DOI: 10.1101/2020.11.02.365031.

Pressey RL, Visconti P, McKinnon MC, Gurney GG, Barnes MD, Glew L, Maron M. 2021. The mismeasure of conservation. Trends in Ecology & Evolution 36:808–821. Elsevier Ltd. Available from https://doi.org/10.1016/j.tree.2021.06.008.

Pynegar EL, Gibbons JM, Asquith NM, Jones JPG. 2019. What role should randomized control trials play in providing the evidence base for conservation? Oryx 55:235–244. Available from https://www.cambridge.org/core/product/identifier/S0030605319000188/type/journal_article.

Rodrigues ASL, Cazalis V. 2020. The multifaceted challenge of evaluating protected area effectiveness. Nature Communications 11:1–4. Springer US. Available from http://dx.doi.org/10.1038/s41467-020-18989-2.

Rose DC et al. 2018. The major barriers to evidence-informed conservation policy and possible solutions. Conservation Letters 11:1–12.

Runting RK, Phinn S, Xie Z, Venter O, Watson JEM. 2020. Opportunities for big data in conservation and sustainability. Nature Communications 11:1–4. Springer US. Available from http://dx.doi.org/10.1038/s41467-020-15870-0.

Salafsky N et al. 2008. A standard lexicon for biodiversity conservation: Unified classifications of threats and actions. Conservation Biology DOI: 10.1111/j.1523-1739.2008.00937.x.

Schleicher J, Eklund J, Barnes M, Geldmann J, Oldekop JA, Jones JPG. 2019. Statistical matching for conservation science. Conservation Biology:cobi.13448. Available from https://onlinelibrary.wiley.com/doi/abs/10.1111/cobi.13448.

Stuart EA. 2010. Matching Methods for Causal Inference: A Review and a Look Forward. Statistical Science 25:1–21.

Terraube J, van doninck J, Helle P, Cabeza M. 2020. Assessing the effectiveness of a national protected area network for carnivore conservation. Nature Communications 11:1–9. Springer US. Available from http://dx.doi.org/10.1038/s41467-020-16792-7.

Tittensor DP et al. 2014. A mid-term analysis of progress toward international biodiversity targets. Science 346:241–244.

Troudet J, Grandcolas P, Blin A, Vignes-Lebbe R, Legendre F. 2017. Taxonomic bias in biodiversity data and societal preferences. Scientific Reports 7:1–14.

UNEP-WCMC, IUCN. 2021. Protected Planet Report 2020. Available from https://livereport.protectedplanet.net/.

Visconti P, Butchart SHM, Brooks TM, Langhammer PF, Marnewick D, Vergara S, Yanosky A, Watson JEM. 2019. Protected area targets post-2020. Science 364:239–242. Available from http://www.sciencemag.org/lookup/doi/10.1126/science.aav6886.

Wauchope H et al. 2019a. Quantifying the impact of protected areas on near-global waterbird population trends, a pre-analysis plan:1–17. Available from https://doi.org/10.7287/peerj.preprints.27741v1.

Wauchope Hannah S, Amano T, Geldmann J, Johnston A, Simmons BI, Sutherland WJ, Jones JPG. 2020. Evaluating impact using time-series data. Trends in Ecology & Evolution In Review:1–10. The Authors. Available from https://doi.org/10.1016/j.tree.2020.11.001.

Wauchope HS, Amano T, Sutherland WJ, Johnston A. 2019b. When can we trust population trends? A method for quantifying the effects of sampling interval and duration. Methods in Ecology and Evolution 10:2067–2078.

Wiik E, Jones JPG, Pynegar E, Bottazzi P, Asquith N, Gibbons J, Kontoleon A. 2020. Mechanisms and impacts of an incentive-based conservation program with evidence from a randomized control trial. Conservation Biology 34:1076–1088.

Williamson EJ, Forbes A. 2014. Introduction to propensity scores. Respirology 19:625–635.

WWF. 2020. Living planet report 2020 - Bending the curve of Biodiversity loss. Page Almond R.E.A., Grooten M. and Petersen, T. (Eds). WWF, Gland, Switzerland. Available from https://f.hubspotusercontent20.net/hubfs/4783129/LPR/PDFs/ENGLISH-FULL.pdf.

