## Supplementary material for "Assessing the global impact of targeted conservation actions on species abundance"

**Supplementary information**

**Taxonomic classes categorized into groups**

We categorized Actinopteri, Coelacanthi, Dipneusti, Elasmobranchii, Holocephali, Myxini and Petromyzonti together as ‘fish’, Aves as ‘birds’, Mammalia as ‘mammals’, Reptilia as ‘reptiles’ and Amphibia as ‘amphibians’.

**Caveats**

The management actions used in this analysis are recorded from information in the LPD and, in few instances, from external sources online. SJ and MH have made every effort to correctly code the actions based on information available in the LPD, original sources and additional online material. However, some errors may remain as management actions have not been coded according to consistent categories. This means that management actions may be in place for some populations but are missing from the LPD (omission error). Also, management actions may be documented in place, but are relatively trivial / inconsequential to the population (commission error) – our analysis treats all actions as equal importance. Furthermore, management information may be documented, but may not reflect all actions in place for that population. Additionally, conservation actions apply only to the population during that time-series time-period.

**Validating the link between conservation actions and reasons for population increase**

For the selected subsample used to estimate the impact of conservation on the global index, we assumed a causal relationship between conservation actions and population increases. However, a potential concern was whether the reasons for population increases listed in the original sources and the conservation actions that we categorized matched. A mismatch between the selected populations’ reasons for population increase and the targeted conservation actions would either mean that (i) the recorded conservation actions did not cause the population increase or (ii), that conservation actions were missing from the original sources or not categorized accordingly. We verified whether the targeted conservation actions and reasons for increase aligned, effectively meaning that conservation was likely caused the population increase, by comparing the frequency at which conservation actions and different reasons for population increase were recorded for populations (Fig S6). If population increases were caused by conservation actions, we would expect a higher frequency at which specific reason for increase logically caused by a certain conservation action and that specific conservation action occurred. For example, we would expect increasing populations targeted by effective legislation to have legal protection listed as the reason for increase. For each population, unique combinations of the selected reasons for population increase and primary conservation type were expanded into separate rows, each row representing a unique combination of a single reason listed as the cause of population increase and a single recorded conservation action. The frequency of unique combinations was then summarized into a frequency table which we used to visualize the frequency at which each of the primary conservation actions occurred relative to each of the selected reasons for increases.

**Table S1** Parameter estimates for the seven primary types of conservation actions, research, utilization status and time series length.

| | Estimate | Std. Error | t value |
|---|---|---|---|
| Utilised | -0.0091880 | 0.0112007 | -0.820 |
| Ts length | 0.0014400 | 0.0006965 | 2.067 |
| Land water protection | 0.2253907 | 0.1165584 | 1.934 |
| Land water management | 0.2003847 | 0.0306416 | 6.540 |
| Species management | 0.1153678 | 0.0254073 | 4.541 |
| Education awareness | -0.0118926 | 0.1442860 | -0.082 |
| Law policy | 0.0209303 | 0.0383110 | 0.546 |
| Incentives | 0.0094938 | 0.1945584 | 0.049 |
| External capacity | 0.4269375 | 0.4397941 | 0.971 |
| Research | -0.2371960 | 0.0762102 | -3.112 |

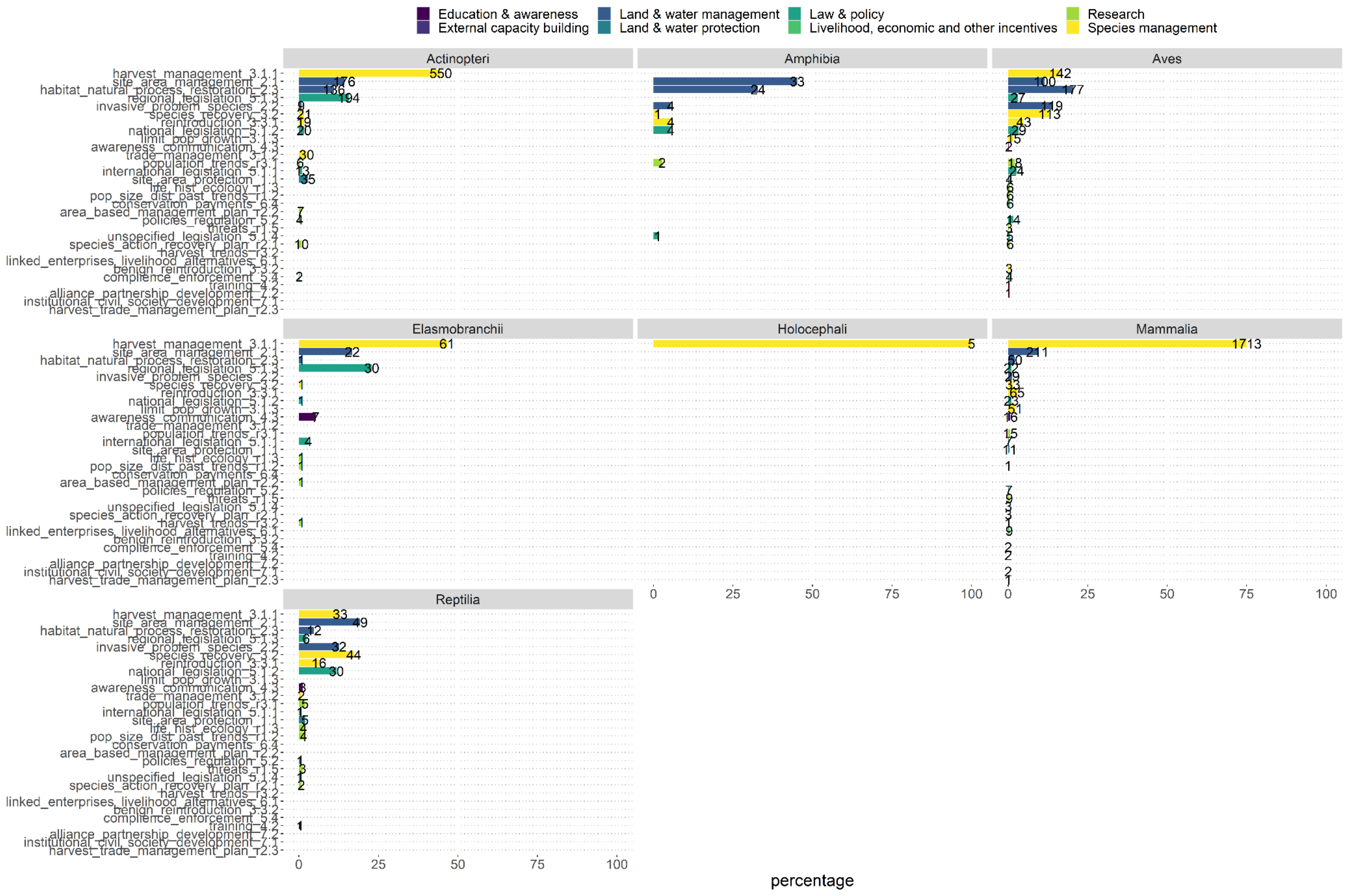

Education & awareness
External capacity building
Land & water management
Land & water protection
Law & policy
Livelihood, economic and other incentives
Research
Species management
Actinopteri
Amphibia
Aves
Elasmobranchii
Holocephali
Mammalia
Reptilia
harvest_management_3.1.1
site_area_management_2.1
habitat_natural_process_restoration_2.3
regional_legislation_5.1.3
invasive_problem_species_2.2
species_recovery_3.2
reintroduction_3.3.1
national_legislation_5.1.2
limit_pop_growth_3.1.3
awareness_communication_4.3
trade_management_3.1.2
population_trends_r3.1
international_legislation_5.1.1
site_area_protection_1.1
life_hist_ecology_r1.3
pop_size_dist_past_trends_r1.2
conservation_payments_6.4
area_based_management_plan_r2.2
policies_regulation_5.2
threats_r1.5
unspecified_legislation_5.1.4
species_action_recovery_plan_r2.1
harvest_trends_r3.2
linked_enterprises_livelihood_alternatives_6.1
benign_reintroduction_3.3.2
compliance_enforcement_5.4
training_4.2
alliance_partnership_development_7.2
institutional_civil_society_development_7.1
harvest_trade_management_plan_r2.3
0
25
50
75
100
percentage

**Figure S1** Number of targeted populations and the relative percentage of detailed conservation actions for taxonomic classes. For each of the taxonomic classes with targeted conservation actions, the x-axis shows the percentage of populations targeted by the detailed conservation actions and research (Salafsky et al. 2008). The number of targeted populations is shown for each bar.

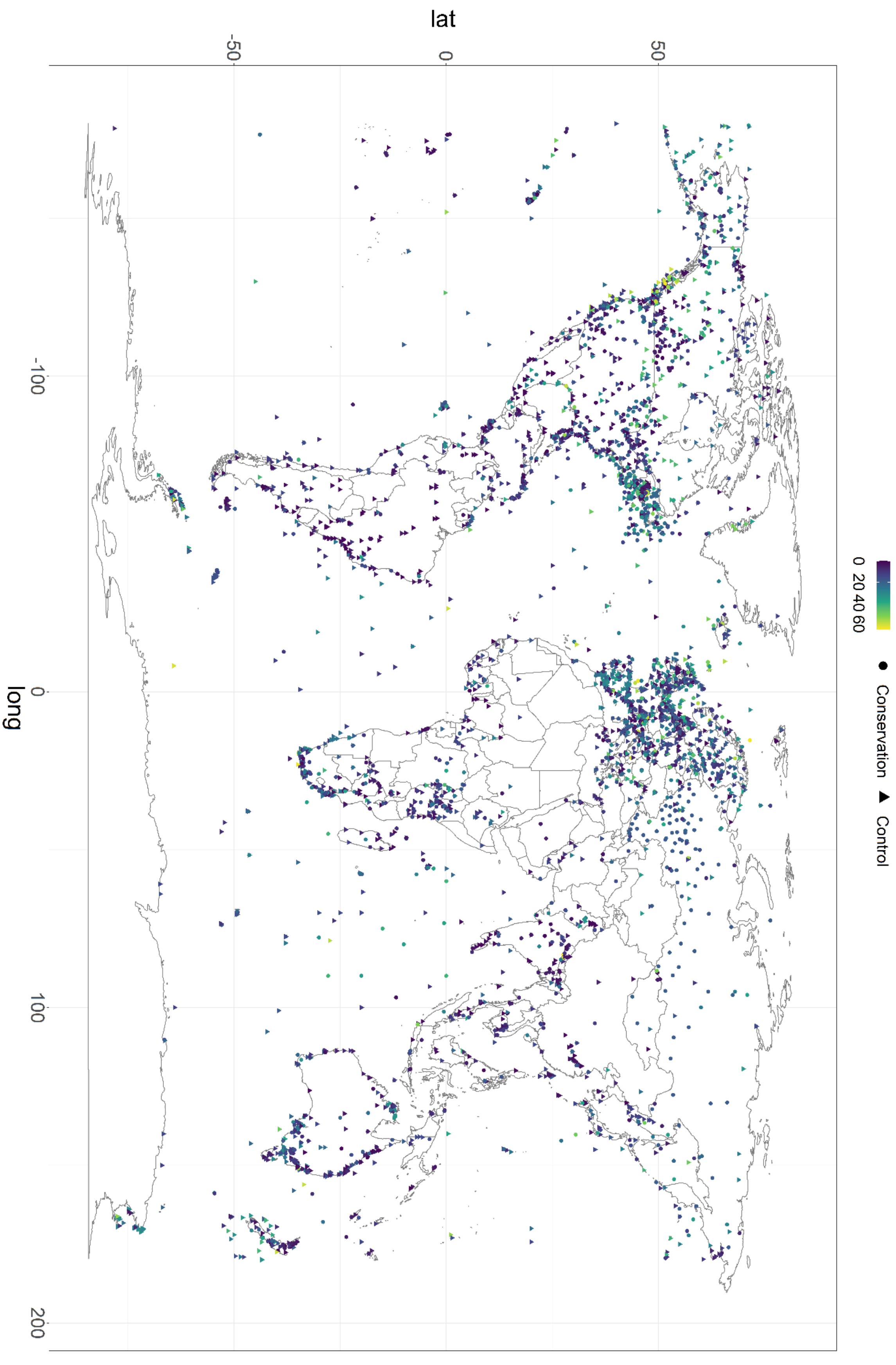


**Figure S2** Length of each managed population.

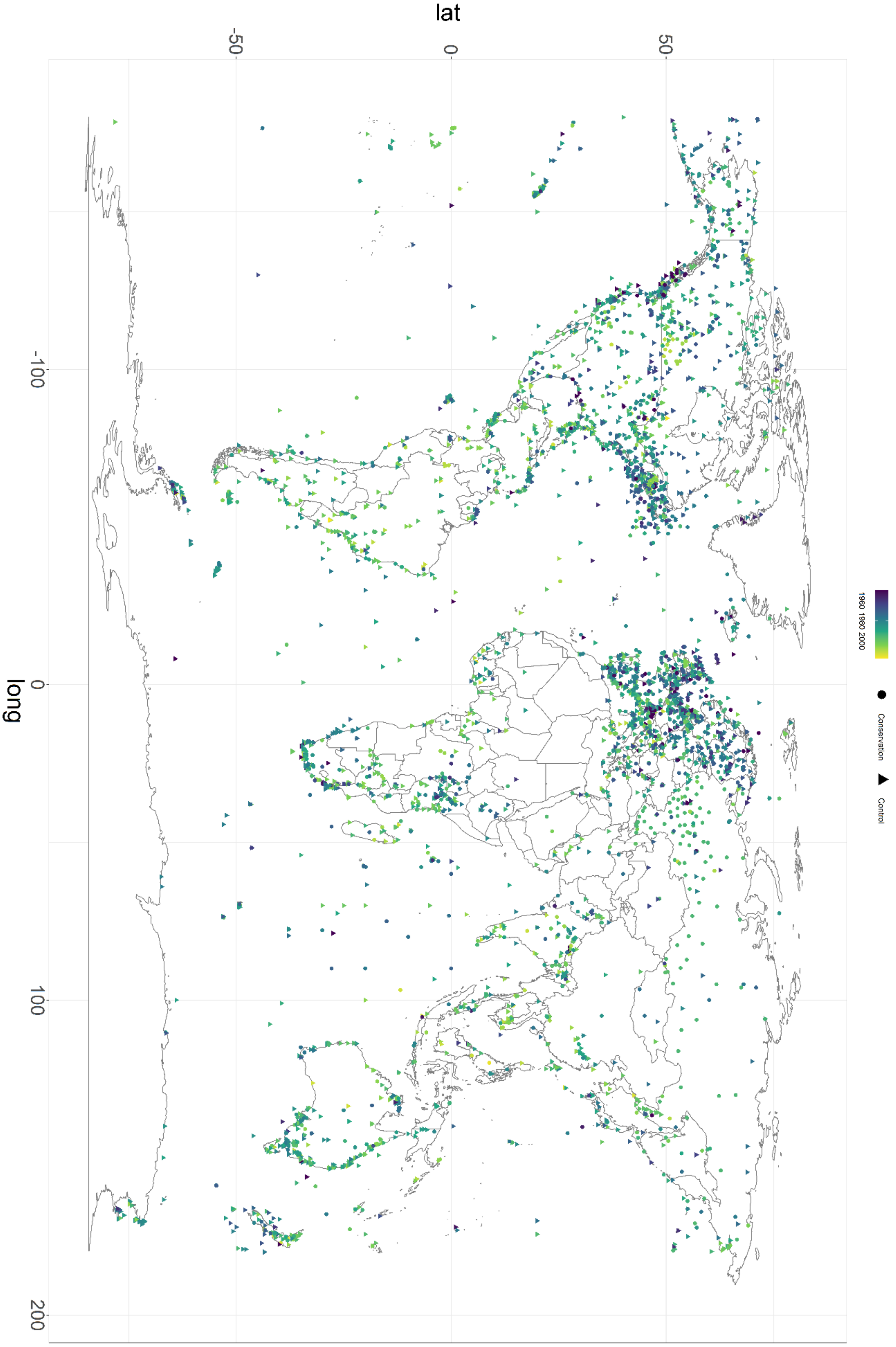


**Figure S3** Starting year of each managed population.

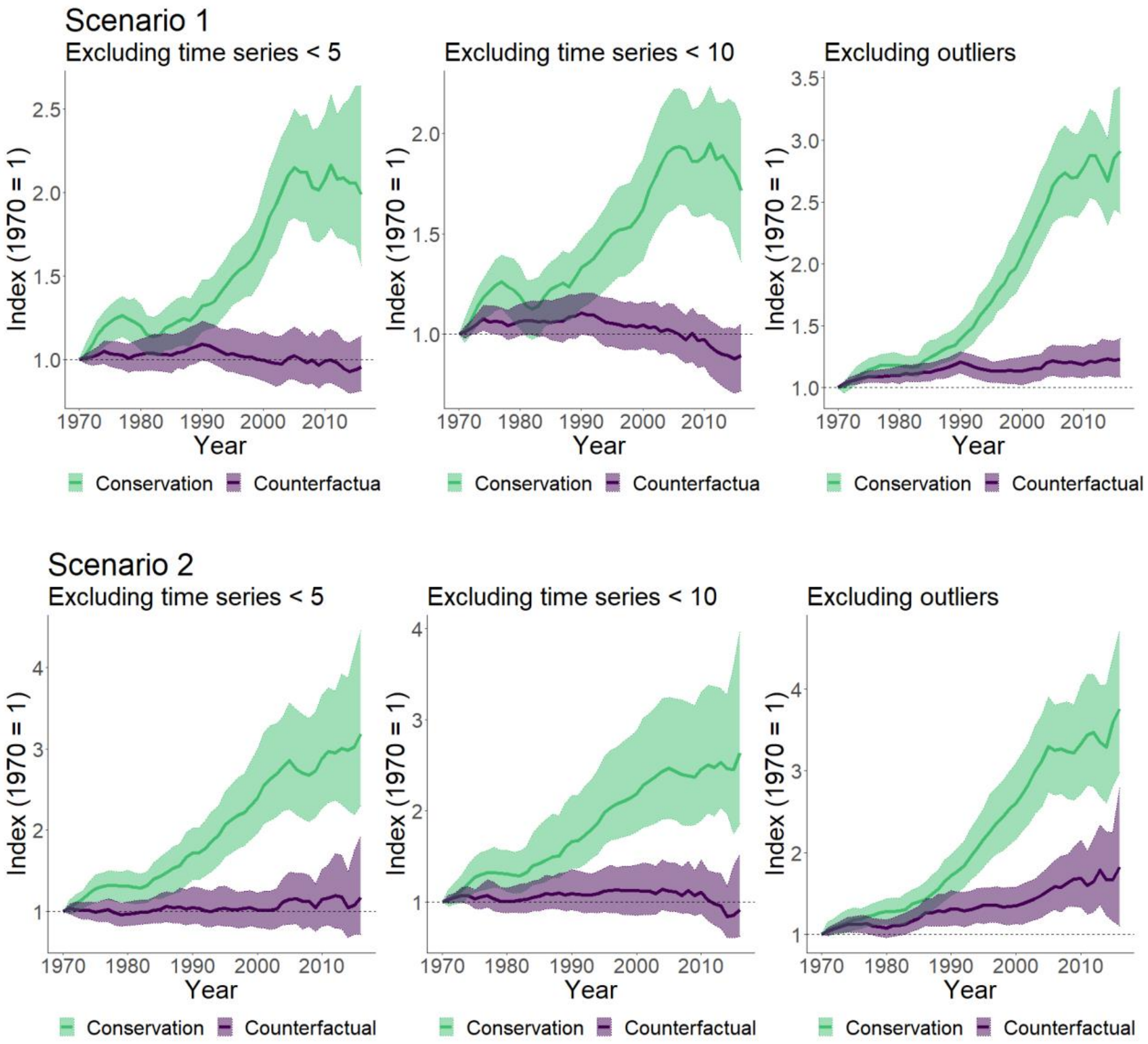

Scenario 1
Excluding time series < 5
Excluding time series < 10
Excluding outliers
Index (1970 = 1)
Year
1970
1980
1990
2000
2010
1.0
1.5
2.0
2.5
3.0
3.5
Conservation
Counterfactua
Conservation
Counterfactua
Conservation
Counterfactual
Scenario 2
Excluding time series < 5
Excluding time series < 10
Excluding outliers
Index (1970 = 1)
Year
1
2
3
4
Conservation
Counterfactual
Conservation
Counterfactual
Conservation
Counterfactual

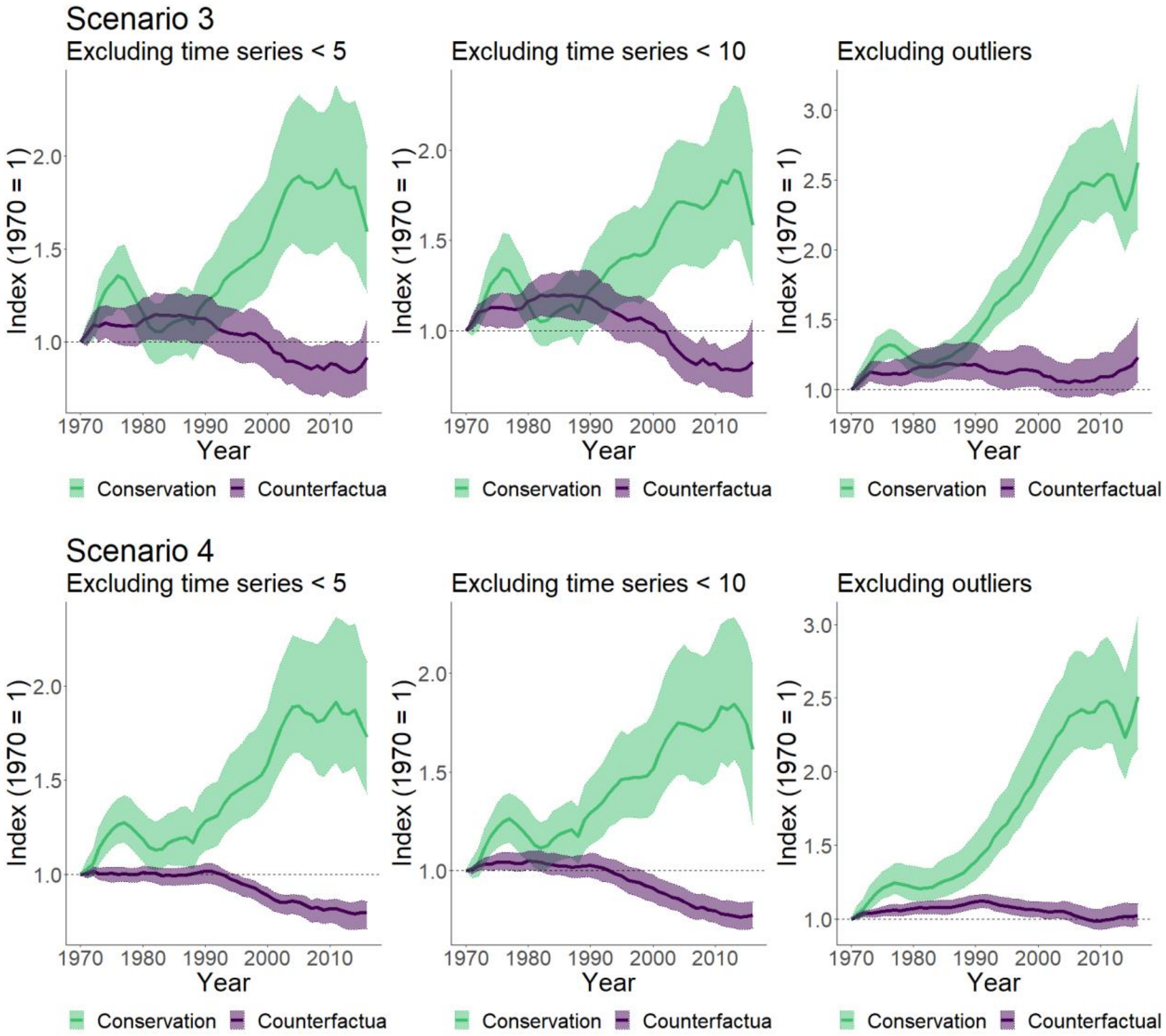


**Figure S4** Sensitivity test for the four scenarios. Trends were calculated excluding populations covering less than five and ten years and excluding species with populations in the top and bottom 1% annual population change quantiles.

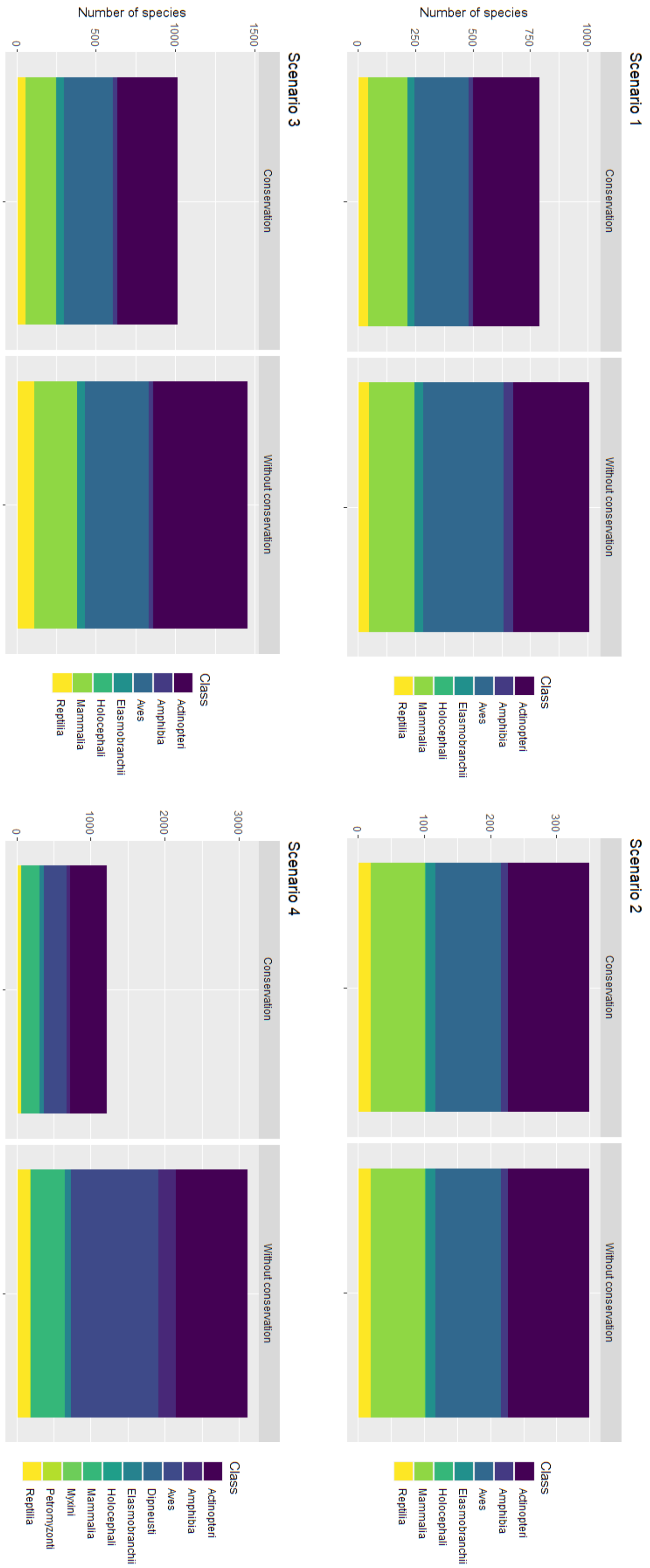


**Figure S5** Number of species within each taxonomical class for each scenario.

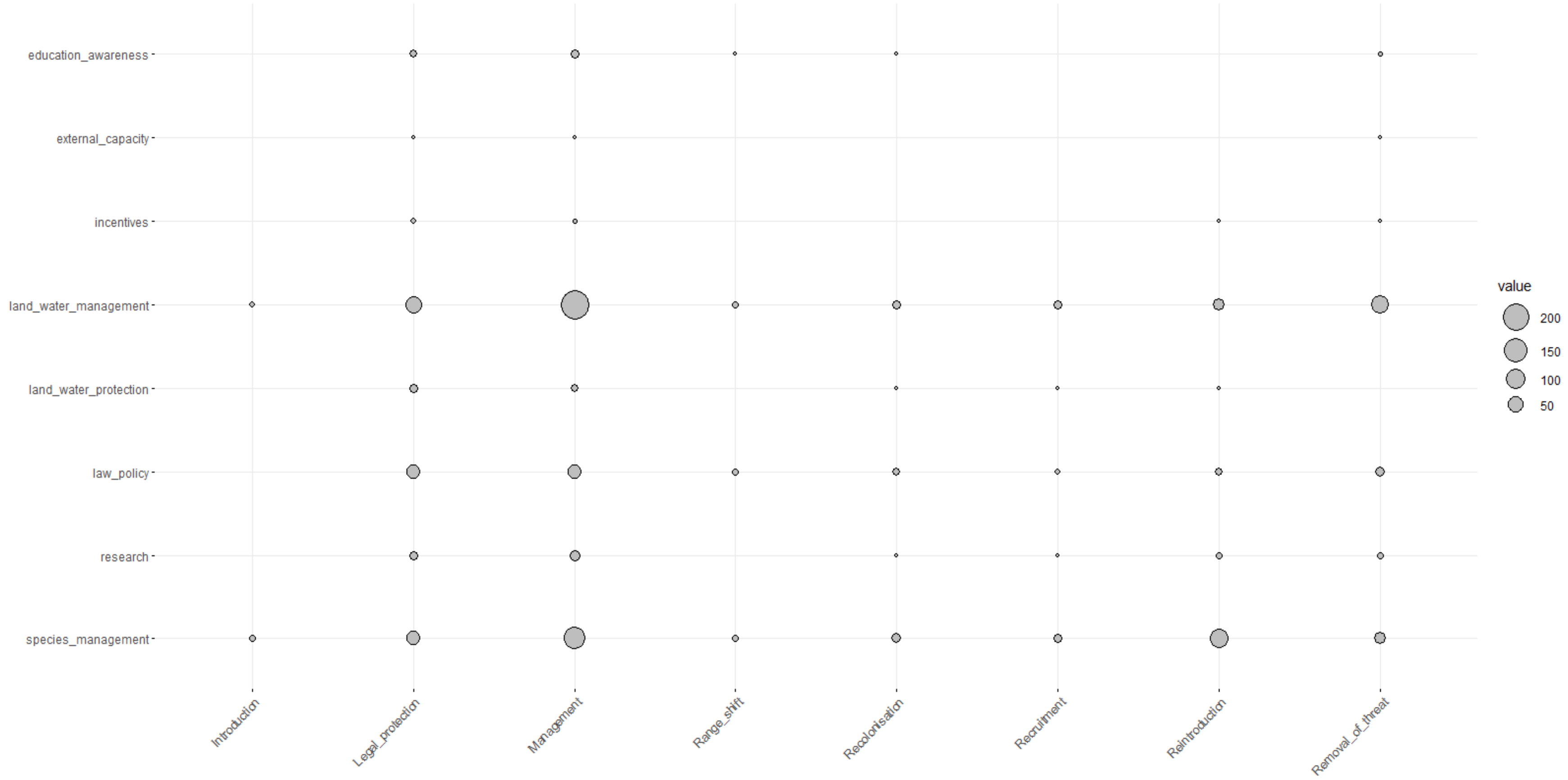
education_awareness
external_capacity
incentives
land_water_management
land_water_protection
law_policy
research
species_management
Introduction
Legal_protection
Management
Range_shift
Recolonisation
Recruitment
Reintroduction
Removal_of_threat
value
200
150
100
50

**Figure S6** Combination frequencies for conservation actions (y-axis) and reasons for increase listed in the LPD (x-axis).

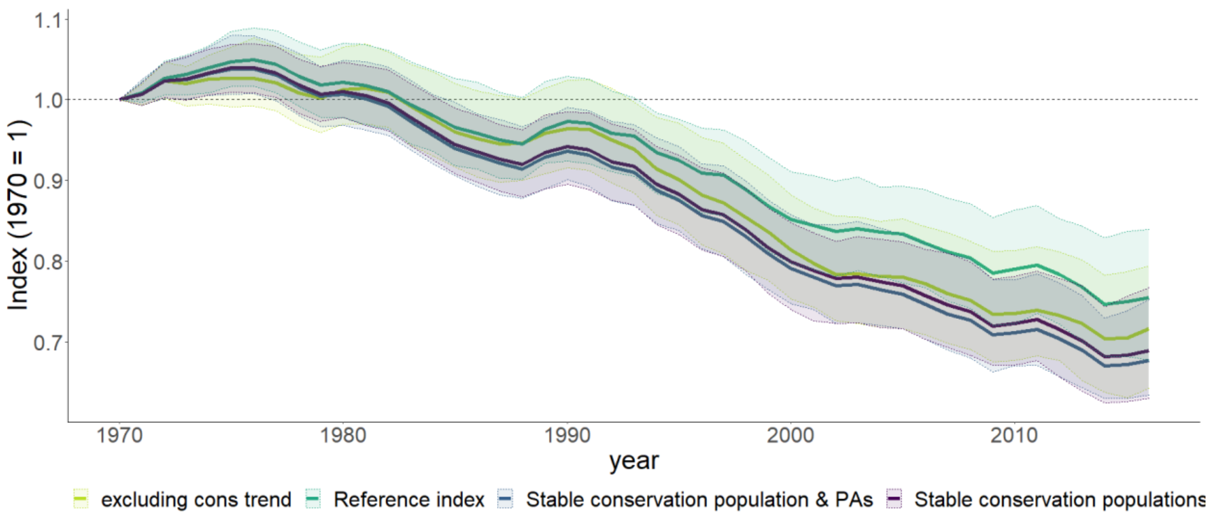


**Figure S7** A reference global vertebrate population index (top dark green line: -24%; nspp = 4,622, npop = 26,871): next, a similar index but excluding all conservation targeted populations (second green line from the top: -28%; nspp = 4226, npop = 21628);  population index assuming stable populations for conservation targeted populations with conservation as the reason for increase (second purple line from the bottom: -31%; nspp = 286, npop = 519): finally, a population index assuming stable populations for conservation targeted populations and populations inside PAs with conservation given as the reason for increase (blue bottom: -32%; nspp = 329, npop = 600). Note that, the number of species and populations listed for the two last indices refers to the number of species and populations assumed stable in the absence of conservation. The number of species and populations used to create the indices are similar to the full reference index.
